# Blockade of TMPRSS2-mediated priming of SARS-CoV-2 by the N-terminal peptide of lactoferrin

**DOI:** 10.1101/2021.12.20.473447

**Authors:** Anna Ohradanova-Repic, Laura Gebetsberger, Gabor Tajti, Gabriela Ondrovičová, Romana Praženicová, Rostislav Skrabana, Peter Baráth, Hannes Stockinger, Vladimir Leksa

**Author notes:** To whom the correspondence should be addressed: Dr. Vladimir Leksa, Institute of Molecular Biology SAS, Dubravska cesta 21, 845 51 Bratislava, Slovakia.

## Abstract

In addition to vaccines, there is an urgent need for supplemental antiviral therapeutics to dampen the persistent COVID-19 pandemic caused by the severe acute respiratory syndrome coronavirus-2 (SARS–CoV-2). The transmembrane protease serine 2 (TMPRSS2), which is responsible for the proteolytic processing of the SARS-CoV-2 spike protein as virus priming for cell entry, appears as a rational therapeutic target for the clearance of SARS-CoV-2 infection. Accordingly, selective inhibitors of TMPRSS2 represent potential tools for prevention and treatment of COVID-19. Here, we tested the inhibitory capacities of the human milk glycoprotein lactoferrin and its N-terminal peptide pLF1, which we identified as inhibitors of plasminogen, a serine protease homologous to TMPRSS2. *In vitro* proteolysis assays revealed that, unlike full-length lactoferrin, pLF1 significantly inhibited the proteolytic activity of TMPRSS2. pLF1 inhibited both the proteolytic processing of the SARS-CoV-2 spike protein and the SARS-CoV-2 infection of simian Vero cells. Because lactoferrin is a natural product and several biologically active peptides, such as the N-terminally derived lactoferricins, are produced naturally by pepsin-mediated digestion, natural or synthetic peptides from lactoferrin represent well-achievable candidates for supporting prevention and treatment of COVID-19.

## Introduction

The global COVID-19 pandemic is unceasingly firing up a demand for cheap and available therapeutics to supplement standard treatment protocols. The virus priming, i.e., the proteolytic processing of the spike (S)-protein of the severe acute respiratory syndrome coronavirus-2 (SARS-CoV-2) by host serine proteases (1,2), in particular the transmembrane protease serine 2 (TMPRSS2) (3), is an event essential for the binding of virus to angiotensin converting enzyme-2 (ACE2), and as consequence, virus entry to the host cell. Thus, the inhibition of virus priming might represent a promising piece in the puzzle of managing the COVID-19 pandemic (4–6). Previously, we identified the human milk glycoprotein lactoferrin (LF) as a potential inhibitor of the serine protease plasminogen (Plg) (7). Here, we expand our previous data to SARS-CoV-2 and show that the very N-terminal peptide of LF inhibits TMPRSS2 and subsequently SARS-CoV-2 priming and infection of target cells.

## Results

Previously, we described the binding of LF to the zymogen Plg resulting in the blocking of conversion of Plg to the serine protease plasmin by urokinase-type plasminogen activator (uPA) (7). According to mapping studies, LF binds to Plg via its highly cationic N-terminal region (7). From this region (residues 1-49), the biologically active natural peptides, called lactoferricins (LFCs), with antiviral and antibacterial activities are derived (8,9). Notably, synthetic analogs of LFCs possess antimicrobial properties similar to natural fragments (9). To specify the role of the N-terminal region in Plg inhibition, we tested the inhibitory capacity of the corresponding synthetic peptide pLF1. As controls, we used three other peptides, pLF2 derived from the helix linker region of LF, pLF3 from the C-terminal part containing a C-terminal lysine, and pCTR, a control peptide, as well as the lysine analogue tranexamic acid (TA), which is known to block the lysine-dependent Plg binding, and the serine proteases’ inhibitor aprotinin (Ap). An *in vitro* activation assay using Plg revealed that LF and pLF1 inhibited Plg activation by uPA (Fig. 1A). However, in contrast to LF, the capability of pLF1 to block Plg activation was uPA-independent because pLF1 was able to inhibit even the intrinsic proteolytic activity of active plasmin with the half maximum concentration (IC50) of 6.9 μg/mL and maximum inhibition was obtained with 20 μg/mL (Fig. 1B). Thus, while LF blocks only the conversion of inactive Plg to the active serine protease plasmin, the N-terminal peptide pLF1 is also able to execute its inhibitory effect directly on active plasmin.

**Figure 1.**
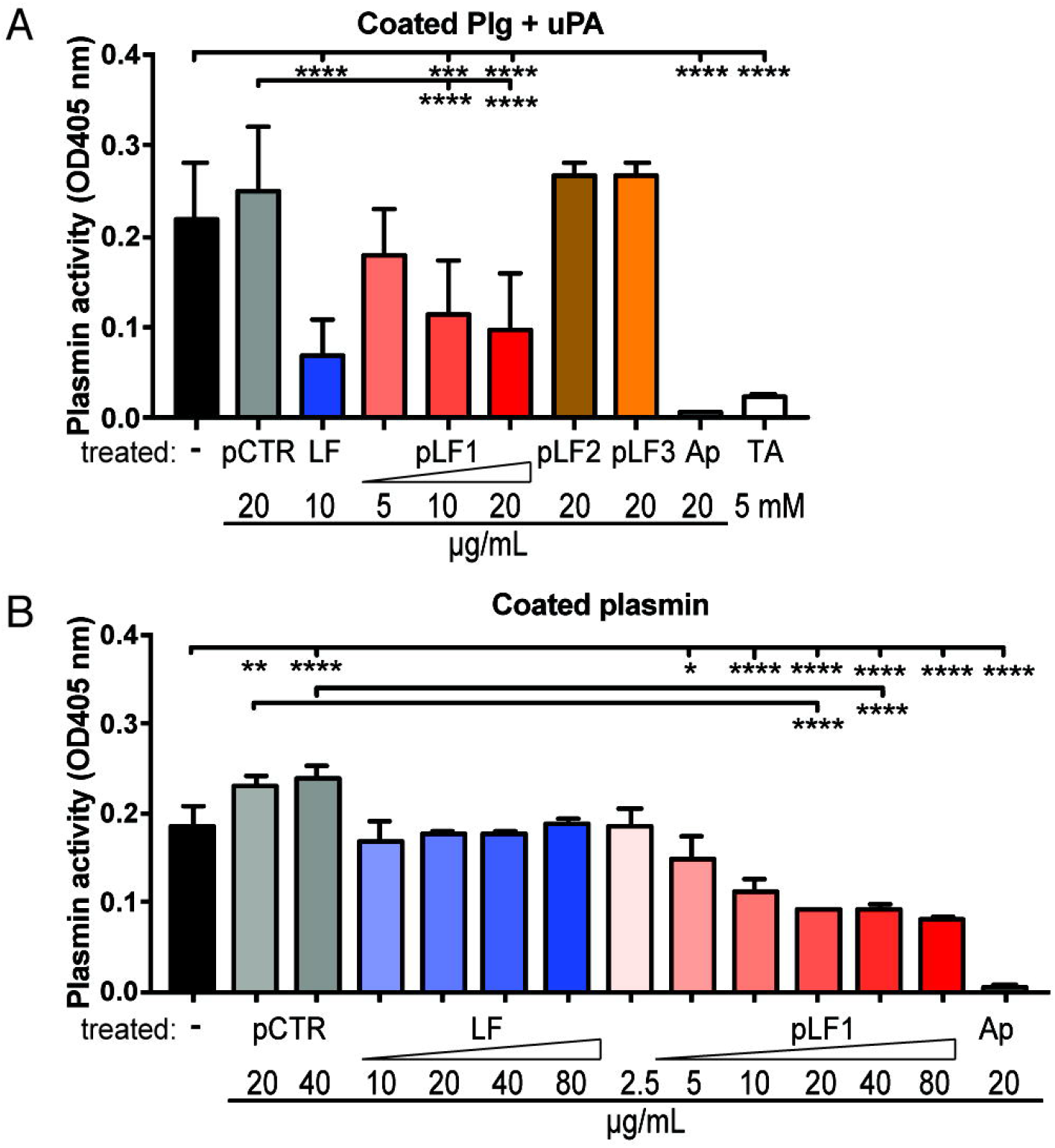
Effect of the LF-derived peptides pLF1-pLF3 on Plg and plasmin activity. *A,* Plg was coated onto wells of a 96-well plate, incubated with either LF, the LF derived peptides pLF1-pLF3, the control peptide (pCTR), Ap (an inhibitor of serine proteases) or TA (a lysine analogue) at the indicated concentrations. Plg was activated by adding uPA at 37°C and its activity was measured by supplementation of the chromogenic plasmin-specific substrate S-2251 using a microplate reader. *B,* Plasmin was coated onto wells, incubated with increasing concentrations of either pCTR, LF or pLF1 as indicated, and supplemented with S-2251. Ap served as positive control. The color change was measured by a microplate reader.

Because pLF1 blocked the intrinsic plasmin activity and the structures of the catalytic domains of serine proteases display a high level of identity (10), we considered that pLF1 might target the catalytic domain directly. Therefore, we tested the inhibitory effect of pLF1 against other serine proteases: namely TMPRSS2, elastase and trypsin. As shown in Fig. 2, indeed pLF1, but neither LF, nor the other tested peptides derived from LF, nor the control peptide pCTR, significantly reduced the proteolytic activity of TMPRSS2 (Fig. 2A), elastase or plasmin (Fig. 2B, C). In contrast, the inhibitory effect of pLF1 on trypsin was not significant (Fig. 2D). pLF1 inhibited TMPRSS2 with IC50 of 9.8 μg/mL with maximum inhibition at 40 μg/mL (Fig. 2A).

**Figure 2.**
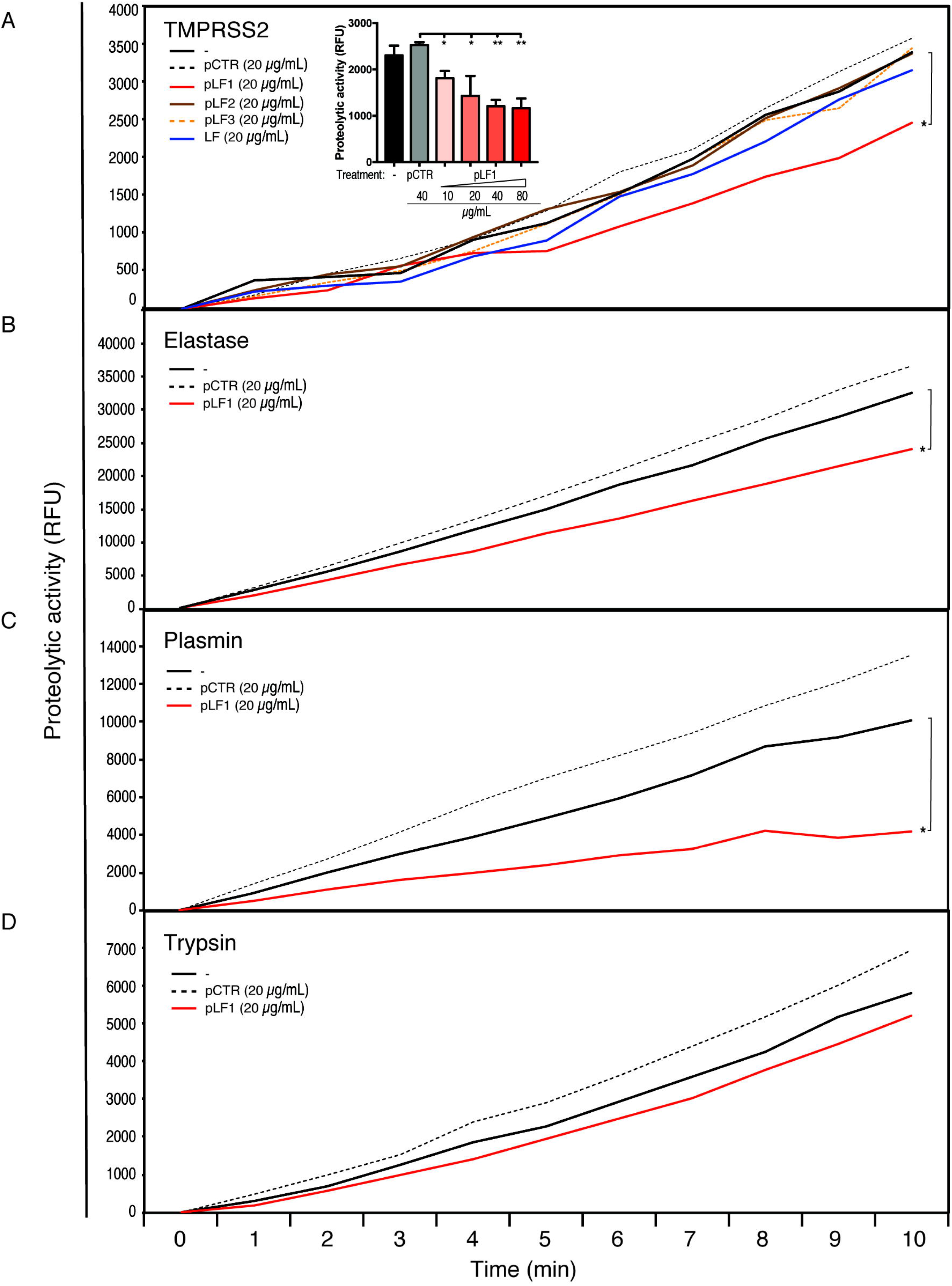
Effect of the LF-derived peptides pLF1-pLF3 on the activity of serine proteases. Proteolytic activities of purified TMPRSS2 (A), elastase (B), plasmin (C), and trypsin (D) were measured with or without peptides pLF1, pLF2, pLF3 or pCTR at 37°C in PBS using the fluorogenic substrate Boc-Gln-Ala-Arg-AMC. The reaction was monitored using an ELISA reader for the indicated time intervals. The *inset* in (A) shows a concentration dependence of the inhibition of the activity of TMPRSS2 by pLF1.

TMPRSS2 was shown to play a central role in the SARS-CoV-2 entry via proteolytic processing of the S-protein of SARS-CoV-2. Because also plasmin, among other serine proteases, was suggested to cleave the SARS-CoV-2 S-protein (1,3,11), we tested whether pLF1 would specifically inhibit the cleavage of the S-protein. Actually, when we exposed the S-protein to TMPRSS2 or plasmin we found blockade of its cleavage in the presence of pLF1, while no such effect was seen with LF (Fig. 3A, B). This experiment demonstrated that the N-terminal fragment of LF inhibited the proteolytic processing of the SARS-CoV-2 S protein by the serine proteases TMPRSS2 and plasmin. However, the key question remained whether pLF1 could also inhibit SARS-CoV-2 infection.

**Figure 3.**
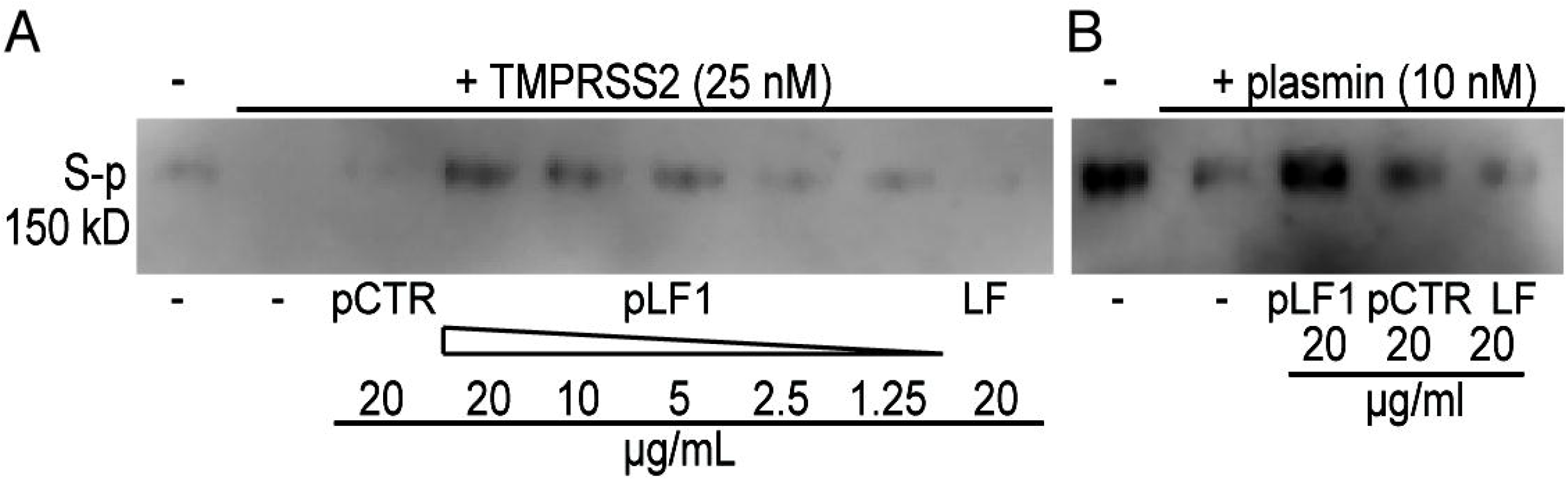
Effect of LF and the LF-derived peptide pLF1 on processing of the SARS-CoV-2 S-protein. Purified recombinant 6xHis-tagged S-protein was exposed to TMPRSS2 (*A*) or plasmin (*B*) with or without LF, pLF1 or pCTR at 37°C in PBS for 30 min. The digestion of the S-protein was analyzed by immunoblotting of the reaction mixtures followed by staining with a SARS-CoV-2 S-protein specific mAb.

To answer this question, we selected Vero cells, which, in contrast to Vero E6 cells, express detectable amounts of TMPRSS2 (12). Before replication-competent SARS-CoV-2 was added, the Vero cells were pre-incubated with LF, the LF-derived peptides pLF1, pLF2, pLF3 or control pCTR, for approximately 40-60 min. As shown in Fig. 4, the N-terminal pLF1 significantly reduced SARS-CoV-2 infection by ≈40% when used at 40 μg/mL, while the other two peptides, pLF2 and pLF3, derived from the middle and C-terminal part of LF, respectively, failed to do so. pLF1 also showed some inhibitory potential at 20 μg/mL (≈30% inhibition), but the differences were not significant. Control peptide pCTR did not show any inhibitory potential. Interestingly, although LF was not able to block TMPRSS2 and plasmin activity (Fig. 2A-B) in the cell-based assay, LF used at 40 μg/mL prevented virus infection by ≈38% (Fig. 4). The latter might be attributed to LF-mediated blockade of cellsurface heparan sulphate proteoglycans that aid coronavirus infection, as published previously (13).

**Figure 4.**
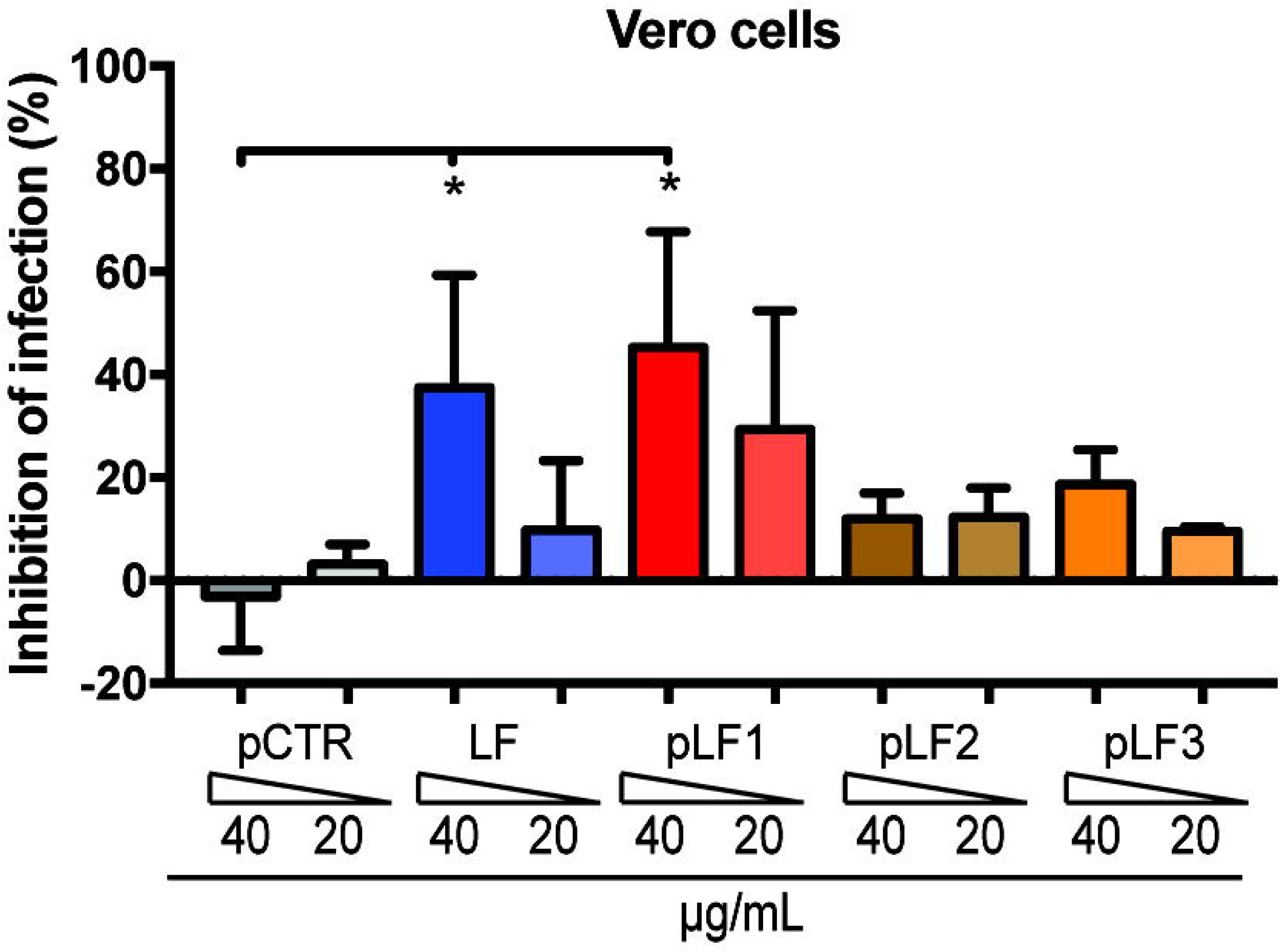
Effect of LF and the LF-derived peptides pLF1-pLF3 on the infection capability of SARS-CoV-2. Vero cells were incubated with or without LF, peptides pLF1, pLF2, pLF3 or pCTR for approx.1 h and then infected with the authentic SARS-CoV-2 (MOI ≈0.02). After 48 h, the cells were fixed and the infection rate was assessed by In-Cell ELISA. Values show mean inhibitory capacity of LF and the indicated peptides ± SD from three independent experiments.

Taken together, our data strongly indicate that the N-terminal peptide fragment of LF inhibits SARS-CoV-2 cell entry through blocking TMPRSS2-mediated virus priming.

## Discussion

As a global threat of the COVID-19 pandemic persists, a request is getting unceasingly stronger for causative therapeutics to complement standard treatment protocols and support a vaccine campaign. The virus priming, essential for the SARS-CoV-2 cell entry via the S-protein – ACE2 interaction, represents a promising therapeutic target. The pharmacological inhibition of TMPRSS2, and also other host proteases that are responsible for processing of the S-protein, is supposed to dampen SARS-CoV-2 cell entry, and hence, its replication (1,2,5,6,14,15).

The human glycoprotein LF is synthetized by exocrine glands and is present at high concentrations in secretory body fluids, particularly in milk, but also within inflammatory granules of neutrophils. LF has been from the beginning of the pandemic recognized as a potential drug candidate (16,17), and recently, it was found to inhibit SARS-CoV-2 infection *in vitro* (18). Moreover, in a recent clinical study a significant effect of the supplementary LF treatment on reduction in COVID-19 symptoms was found (19). This effect was mainly attributed to the previously characterized ability of LF to prevent virus binding to cell-surface heparan sulphate proteoglycans on host cells (13), and also to its immunomodulatory properties (20).

In this report, we demonstrate that LF may contribute to the defense against SARS-CoV-2 infection, in particular via action of its natural N-terminal fragment. Several biologically active natural peptides derived by pepsin cleavage from the N-terminus of both bovine and human LF have been described and termed lactoferricins (LFCs) (9,21,22). LFCs possess antibacterial and antiviral activities (23) and antiviral effects of LF-derived peptides against SARS-CoV-2 were predicted by simulation (24). Here, we show that the very N-terminal peptide of LF, pLF1, inhibits S-protein priming and infection of SARS-CoV-2.

Previously we showed that LF binds through its N-terminal domain encompassing the LFCs region to the serine protease Plg, and blocks Plg activation by uPA (7). In contrast, pLF1, inhibits Plg activity via a different manner. It blocks directly the intrinsic activity of active plasmin and TMPRSS2 with IC50 6.9 μg/mL and 9.8 μg/mL, respectively.

Representative data on LFCs’ levels in human sera are not available, however, since LFCs are digestion product derived from LF, data on serum LF levels might be correlative. According to our unpublished measurements and also published studies of others, LF levels in blood sera of healthy donors may vary within a dynamic range up to μg/mL values (25), which makes the effective concentrations around IC50 achievable.

Taken together, our results reveal that not only LF but also its bioactive digestion products may be protective against COVID-19, however, via different mechanisms, which can produce synergic therapeutic or prophylactic effects. LF is present at high concentrations in milk, but also in other dairy products and food supplements. Orally administered LF endows a major clinical benefit in human, particularly neonatal medicine (26), and it is considered safe and without adverse effects (27). This makes LF a cheap and widely available candidate for supplementary therapy in management of COVID-19.

## Experimental procedures

### Peptides

The peptides derived from LF were synthesized by Peptide 2.0 (Chantilly, VA). The sequences of the 19-amino-acids-long synthetic peptides were as follows: GRRRSVQWCAVSQPEATKC (the N-terminal pLF1, residues 1-19), EDAIWNLLRQAQEKFGKDK (the middle pLF2, residues 264-282), NLKKCSTSPLLEACEFLRK (the C-terminal pLF3, residues 674-692), and NFRTKSCPLELAKELKLCS (pCTR, a scrambled variant of pLF3).

### In vitro proteolysis assay

For the analysis of Plg activation, uPA (10 nM, Technoclone, Vienna, Austria) was coated in PBS on a 96-well Falcon transparent plate for 2 h at 37°C. The wells were blocked with 1% BSA, washed, and then incubated in PBS at 37°C with human Glu-Plg (50 nM, CoaChrom Diagnostica, Vienna, Austria), inhibitory and control compounds, and the chromogenic plasmin substrate S-2251 (0.8 mM, CoaChrom Diagnostica). Optionally, the wells were coated directly with the purified active plasmin (10 mU/mL, Sigma-Aldrich, St. Louis, MO).

The proteolytic activities of trypsin (10 nM), plasmin (10 nM), elastase (20 nM; all three from Sigma-Aldrich) and TMPRSS2 (25 nM; MyBioSource, San Diego, CA) were measured in a Nunc black fluorescence-based cell assay plate (Sigma-Aldrich) in Tris-HCl buffer (50 mM, pH 8.0, 150 mM NaCl). The proteases were incubated with or without inhibitory and control compounds at 37°C for 30 min, and then the fluorogenic substrate (Boc-Gln-Ala-Arg-AMC; 25 μM; ENZO Life Sciences, Lörrach, Germany) was added. The reaction was measured using an ELISA plate reader (Synergy H1 BioTek microplate reader, Winooski, Vermont, U.S.) under an excitation wavelength of 380 nm and an emission wavelength of 460 nm at the indicated time intervals.

In the SARS-CoV-2 S-protein digestion assay, purified recombinant 6xHis-tagged S-protein (1 μg; Native Antigen, Kidlington, UK) was exposed to TMPRSS2 or plasmin (25 nM or 10 nM, respectively) with or without inhibitory and control compounds for 60 min at 37°C. The reaction mixtures were analyzed by immunoblotting with a specific anti-S-protein monoclonal antibody (mAb) (NovusBio, Littleton, CO).

### Immunoblotting

Samples obtained from *in vitro* proteolysis assay were analyzed by electrophoresis on an 8% SDS-polyacrylamide gel followed by transfer to Immobilon polyvinylidene difluoride membranes (Millipore Co., Bedford, MA). The membranes were blocked using 4% non-fat milk and immunostained with specific mAb. For visualization of proteins, the chemiluminescence image analyzer Azure 280 (AzureBiosystems, Dublin, CA) was used.

### SARS-CoV-2 infection and In-Cell ELISA

The African green monkey kidney-derived Vero cells (ATCC CCL-81^TM^) were seeded into 96-well flat-bottom plates (1×10^4^ cells, 80 μl/well) in Dulbecco’s Modified Eagle’s medium (DMEM), high glucose, with GlutaMAX and sodium pyruvate (Gibco/Thermo Fisher Scientific, Waltham, MA USA) supplemented with 10% fetal calf serum (FCS, Biowest, Nuaillé, France), 1% MEM non-essential amino acids solution, 100 U/mL penicillin and 100 μg/mL streptomycin (all latter from Gibco/Thermo Fisher Scientific). On the next day, tested peptides, LF and controls were diluted in DMEM medium with reduced serum (2% FCS) in a separate plate so that the concentration was 1.33-fold higher than final. Ninety μl of the solution was used to pretreat seeded cells (old medium was discarded) for approx. 40-60 min. During the incubation time, cells were transferred to the BSL3 facility of the Medical University of Vienna and then infected with 30 μl (300 TCID50/well; multiplicity of infection (MOI) ≈0.02) of the authentic SARS-CoV-2 isolate BetaCoV/Munich/BavPat1/2020 [kindly provided by Christian Drosten, Charité, Berlin (28), and distributed by the European Virology Archive (Ref-SKU: 026V-03883)] that was diluted to 1×10^4^ TCID50/mL in DMEM/2% FCS. Cells were then incubated at 37°C. After 48 h, cells were fixed by adding 45 ul/well 37% formaldehyde for 20 min. Supernatants were then aspired and the entire plate was fixed in fresh 5% formaldehyde in PBS for 30 min. Next, the formaldehyde was removed, the cells were washed with PBS, permeabilized using 0.1% Triton X-100 in PBS and blocked with blocking buffer (10% FCS in PBS+0.05% Tween-20). Subsequently, the cells were stained by indirect immunofluorescence using a rabbit anti-SARS-CoV-2 nucleocapsid mAb (40143-R019, SinoBiological, Beijing, China, diluted 1:15000 in blocking buffer) followed by a goat anti-rabbit-HRP conjugate (170-6515, Bio-Rad, Hercules, CA USA, diluted 1:10,000 in blocking buffer). In-Cell ELISA was then developed using DY999 substrate solution according to the manufacturer’s recommendations (R&D Systems, Minneapolis, MN USA) and measured at 450 nm (and at 630 nm to assess the background) using a Mithras multimode plate reader (Berthold Technologies, Bad Wildbad, Germany). To calculate percent inhibition of infection at each well, the following formula was used: 100 – [(X – average of □no virus□ ≥ wells)/(average of ≥ □virus only□ wells - average of □no virus□ wells)*100], where X is the background-subtracted read for each well.

### Statistical analysis

All experiments were performed at least three times in at least duplicates. The data were expressed as mean values with standard deviations. Statistical significance was evaluated by using a Student’s t-test or one-way ANOVA with Tukey post-test; values of p*<0.05, p**<0.005, p***<0.0005 (as indicated) were considered to be significant or highly significant, respectively. Relative IC50 values were determined by a 4PL nonlinear regression curve fit.

## Acknowledgement

This work was supported by grants of the Science and Technology Assistance Agency of the Slovak Republic (APVV-16-0452, APVV-20-0513), of the Slovak Grant Agency VEGA (2/0020/17, 2/0152/21) and of the Austrian Science Fund (FWF; P 34253-B).

## Author information

The authors declare no competing financial interests.

## Authorship contributions

V.L. and A.O.R. conceived and designed the experiments. V.L., A.O.R., L.G., G.T., R.P., R.S., P.B., G.O. performed and analyzed the experiments. H.S. contributed with ideas, comments and materials. V.L. wrote the paper. All authors read and corrected the manuscript.

## Abbreviations

ACE2: angiotensin converting enzyme-2
Ap: aprotinin
COVID-19: coronavirus disease 2019
LF: lactoferrin
LFC: lactoferricin
mAb: monoclonal antibody
Plg: plasminogen
SAR-CoV-2: severe acute respiratory syndrome coronavirus 2
TA: tranexamic acid
TMPRSS2: transmembrane protease serine 2
uPA: urokinase-type plasminogen activator

## Notes

### Competing Interest Statement

The authors have declared no competing interest.

